# Neural tracking of melodic prediction is pre-attentive

**DOI:** 10.1101/2025.10.20.683459

**Authors:** Mika Nash, Magdalena Kachlicka, Marcus Pearce, Pubudu Pathirana, Adam Tierney

## Abstract

Music’s ability to modulate arousal and manipulate emotions relies upon formation and violation of predictions. Music is often used to modulate arousal and mood while individuals focus on other tasks, suggesting that sophisticated musical prediction may be implicit and obligatory, operating regardless of attention. Here we investigated the role of attention in musical prediction by presenting participants simultaneously with a musical passage in one ear and an audiobook in the other. Participants were asked to attend to either speech or music to perform a recognition memory task. Musical prediction was investigated using a computational model of music which was trained on a corpus of Western music (long-term learning) and/or incrementally on each stimulus (short-term learning). While neural tracking of the acoustics of music, including encoding of note onset and pitch interval size, was enhanced by attention, musical prediction tracked the neural signal robustly regardless of attention. However, although neural tracking of long-term musical prediction based on previously-learned statistical patterns was unaffected by attention, tracking of short-term musical prediction learned from the structure of each stimulus was present only when the music was attended. Thus, attention is crucial for short-term musical learning and prediction but is not required for musical prediction based on long-term stylistic exposure.

## Introduction

Much of our perception of the world takes place outside of the focus of attention. This is perhaps especially true of auditory perception, which is less constrained by body position with respect to the environment than other senses. Listening to music, for example, can be a profound aesthetic experience, but in daily life, listeners often put music on and then direct attention to other tasks (1-2). In film a musical score can colour a scene’s emotional impact (3) and enhance memory for plot details (4), but the music is often not attended or remembered (3). Music’s common role as a supporting art form suggests that it can be unconsciously processed, but it remains an open question which features of music can be tracked pre-attentively and which require attention. Understanding the extent to which different musical features can be processed when attention is directed elsewhere would help reveal the characteristics and limitations of pre-attentive processing. Understanding the role of attention in music listening could also help musicians decide what aspects of music to focus on when composing music which is designed to play a supporting role, as is often the case in film and television.

As people listen to music their perceptual systems make predictions: when the next note will arrive, what pitch it will have, and what chord will be played next. Computational models explain how listeners make rhythmic (5-6), melodic (7), and harmonic (8-9) predictions (10). These models analyze musical sequences across an entire corpus and then apply the resulting knowledge to predict features of a musical piece as it unfolds. Musical events with unexpected features, as defined by these models, are linked to lower expectedness ratings (11), longer reaction times (9), larger electrophysiological (EEG) responses (12), greater arousal (13), and characteristic changes in pleasure (14), confirming that prediction is a core component of music listening. However, it remains unknown whether attention is necessary for listeners to form predictions of upcoming musical events based on prior context.

Musical prediction can be understood as an instance of predictive coding, in which individuals form generative models about underlying causes of perceptual data (15). Predictions generated by these models are compared to incoming perceptual data, and prediction errors are used to fine-tune the models. According to predictive coding theory, attention up-weights prediction errors, leading to greater model adjustments (16). Empirical evidence supports this idea that attention can enhance neural responses to failed predictions. For example, some research has found that attention enhances processing of simple unpredictable stimuli such as tones and visual gratings (17-19), while neural responses to predictable stimuli are unaffected by attention. Attention may be particularly necessary for complex predictions made during ecologically valid perception. Brodbeck et al. (20), for example, found that phonemic entropy (i.e. the precision of phonemic predictions) was only a significant predictor of the MEG signal when speech was attended. Prior research and theory, therefore, suggests that attention modulates prediction, and may even be necessary for complex higher-level prediction to take place. However, music’s tendency to fade into the background suggests that it may be designed to be pre-attentively processed, which may limit the role of attention in modulating perception of music. Music has the unique ability to use surprisal to induce pleasure even when highly familiar (21), suggesting that musical expectations may be implicit and obligatory, not requiring attention (22-23).

Here we examined the extent to which musical prediction is modulated by attention. To answer this question, we made use of an EEG paradigm in which speech is played to one ear and music to the other. Participants were asked to attend to either speech or music to perform a recognition memory task. We investigated whether attention modulated basic acoustic features of music, including note onset and pitch, as well as musical prediction. Measures of musical prediction included pitch interval size (since larger pitch intervals violate the statistical pattern of predominance of small intervals in music (11)) and melodic surprisal, as derived from a computational model of auditory expectation (7). Prior research has established that melodic surprisal is tracked by the EEG signal (24), but it remains unknown whether neural tracking of melodic surprisal is modulated by attention. We examined musical prediction generated by a long-term model trained on a corpus of Western music, as well as a short-term model trained incrementally on each stimulus. This enabled us to separately investigate the effect of attention on musical prediction (long-term model) as well as statistical learning (short-term model).

We also asked whether attention modulated the neural tracking of musical beats. Time periods which align with the beat are more likely to feature note onsets, and so incorporating beat structure into computational models improves prediction of note timing (5-6). According to predictive coding, the beat can be seen as a generative model, with off-beat notes leading to prediction error (25-26). Reaction times are faster for stimuli that are aligned with musical beats (27), showing that listeners use beat structure to predict stimulus timing. Prior research on effects of attention on musical beat tracking has shown conflicting results, with some studies reporting that musical beats are represented in the brain even when attention is directed elsewhere (28), while other studies suggest that attention is required for beat perception (29). However, these studies have used simple rhythmic patterns; it remains unclear, therefore, how attention modulates rhythmic prediction during ecologically valid music listening. We assessed the extent to which beat perception draws on attentional resources by using frequency tagging (30) to compare neural tracking of the beat when the music was attended versus ignored.

Finally, a secondary goal of this research was to investigate whether musical training modulates the effects of attention on musical prediction. Musicians are better able to hear music in noise (31) and show better neural tracking of the amplitude contour of naturalistic music (32). However, prior research on effects of musical expertise on musical prediction error responses has found inconsistent results (33-34), and it remains unknown how expertise modulates the effects of attention on prediction error. We asked whether the effect of attention on musical prediction error responses is greater in participants with more years of musical training.

## Results

Performance on the attention task was overall neither at floor nor at ceiling (mean performance = 83.4%, standard deviation = 12.2%). This confirms that the recognition memory task was difficult enough to require focus yet easy enough to be accessible to most of the participants. (As described in the Participants section, two participants needed to be removed due to performing below 50%; these data points are not included in the summary of performance above.)

Neural tracking of speech amplitude was greater when the speech was attended (mean = 0.034, S.D. = 0.014) than when it was ignored (mean = 0.026, S.D. = 0.014; F = 28.5, p < 0.001). This finding, combined with the performance data, demonstrates that our participants were successfully switching attention between the two sound streams. Speech amplitude tracking was also greater in older participants (F = 12.3, p = 0.001), but was not modulated by degree of musical training (p > 0.05). Figure 1 displays neural tracking of speech amplitude across conditions.

**Figure 1.**
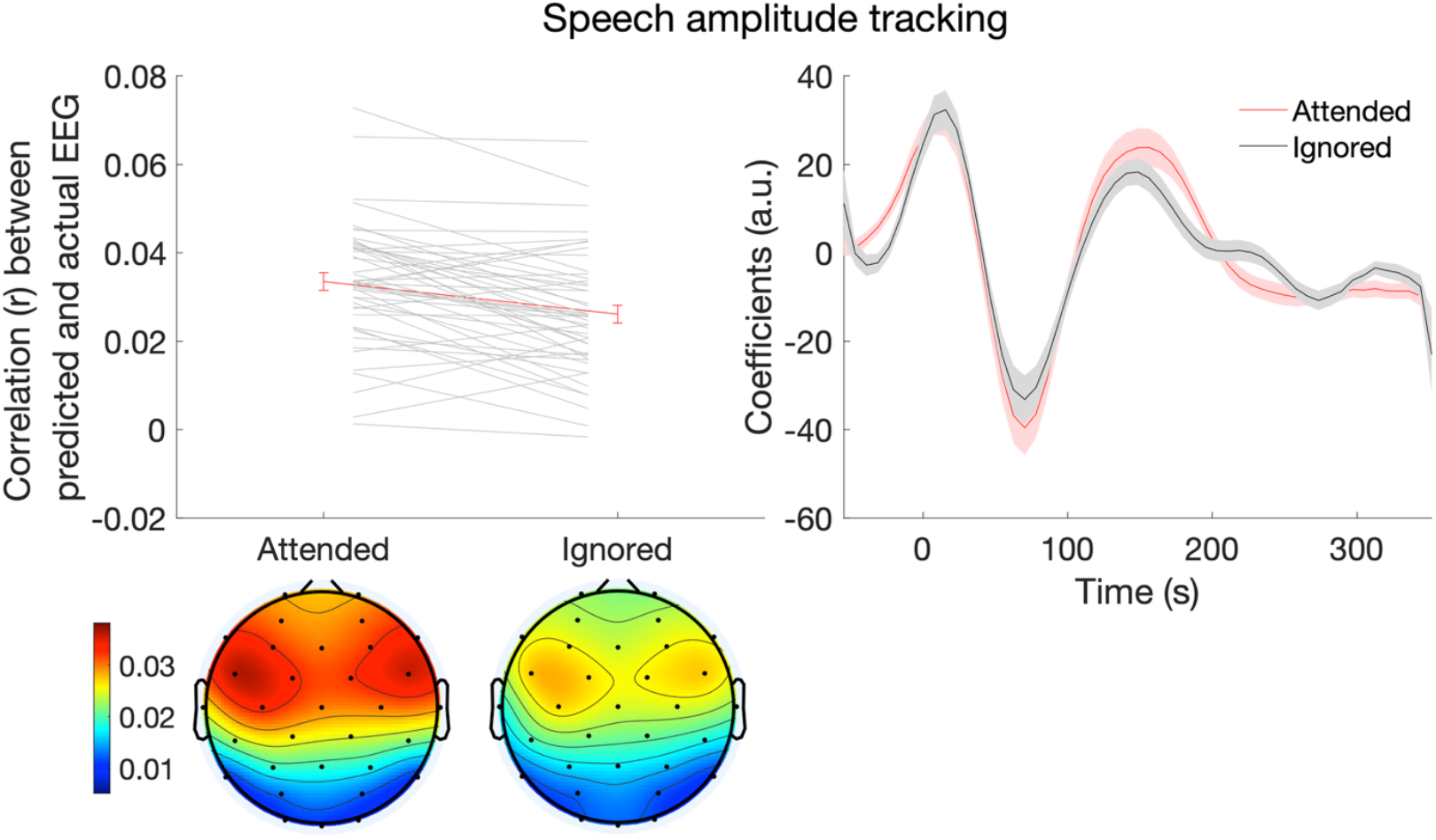
Effect of selective attention on speech amplitude tracking. Attention was directed either towards the speech or a competing stream of music, so that participants could complete a recognition memory task after each trial. The outcome measure plotted on the left is the extent to which changes in the EEG can be predicted by a statistical model making use of speech amplitude as a predictor, with leave-one-trial-out cross-validation. The right-hand side of the figure plots model coefficients across participants. Error bars display one standard error of the mean.

Neural tracking of several characteristics of music was modulated by attention. First, neural tracking of musical note onsets was greater when music was attended (mean = 0.057, S.D. = 0.026) than when it was unattended (mean = 0.050, S.D = 0.020; F = 9.1, p = 0.004). Moreover, musical note onset tracking was greater in older participants (F = 4.9, p = 0.03). In addition, although musical training was not a significant predictor of speech amplitude tracking (p > 0.05), it was a significant predictor of musical note onset tracking (F = 12.4, p = 0.001). Figure 2 displays neural tracking of musical note onset across conditions. Neural tracking of pitch interval size was also greater when music was attended (mean = 0.010, S.D. = 0.0092) versus when it was ignored (mean = 0.0060, S.D. = 0.0095; F = 6.23, p = 0.016). There were no other significant predictors of pitch interval size tracking. Figure 3 displays neural tracking of pitch interval size across conditions.

**Figure 2.**
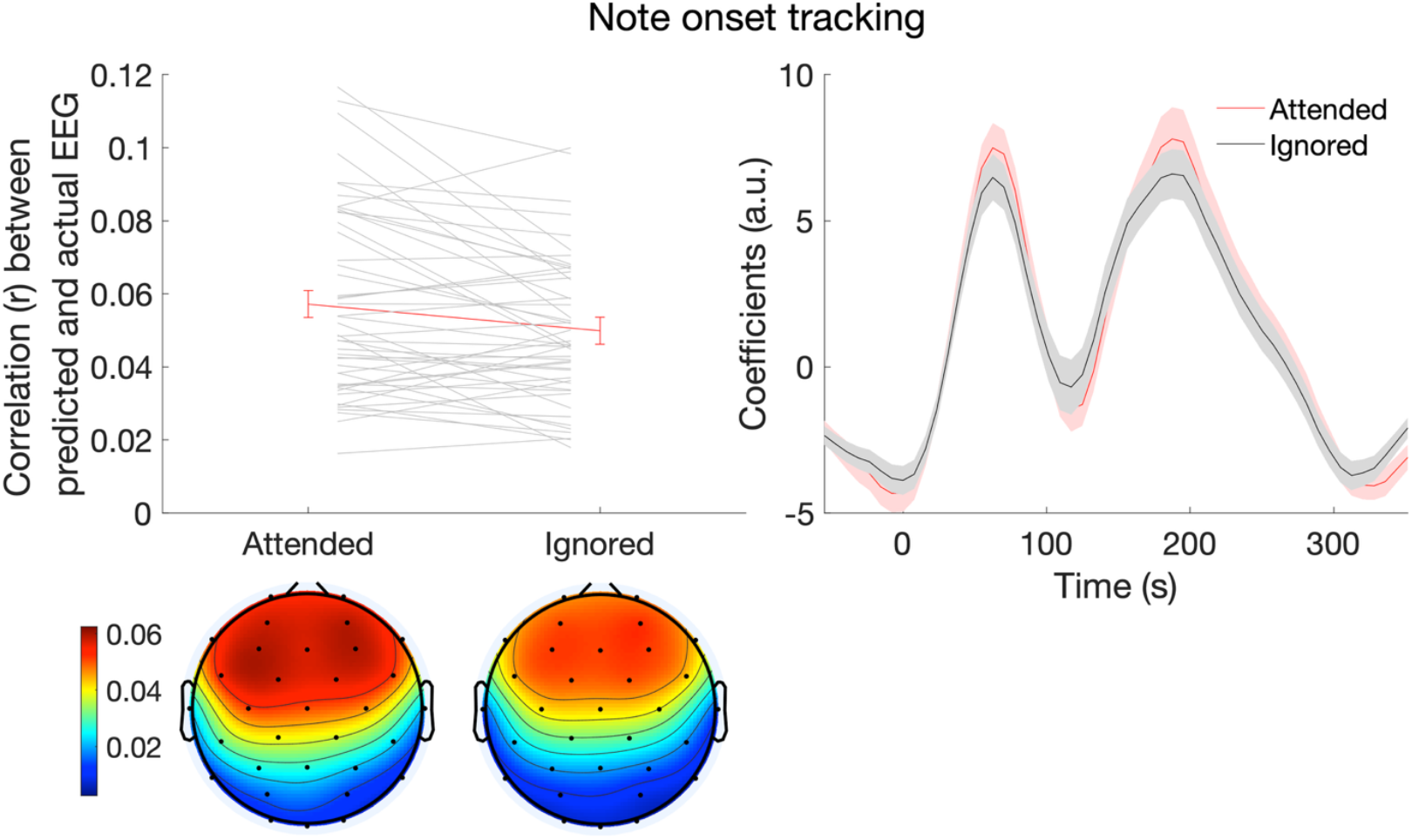
Effect of selective attention on musical note onset tracking. The outcome measure plotted on the left is the extent to which changes in the EEG can be predicted by a model making use of note onset as a predictor. The right-hand side of the figure plots model coefficients across participants. Error bars display one standard error of the mean.

**Figure 3.**
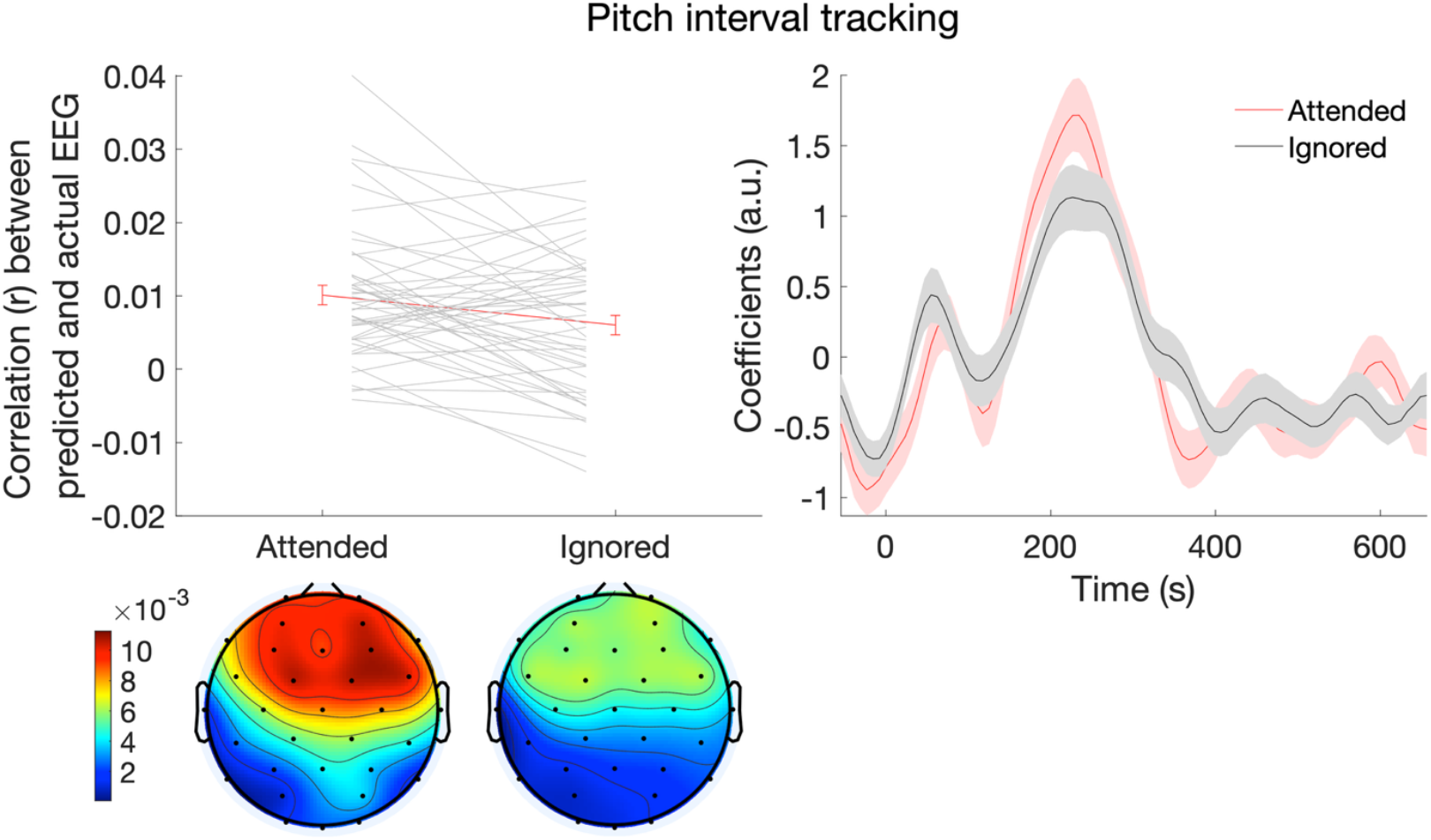
Effect of selective attention on musical pitch interval tracking. The outcome measure plotted on the left is the extent to which changes in the EEG can be predicted by a model making use of absolute pitch interval size as a predictor. The right-hand side of the figure plots model coefficients across participants. Error bars display one standard error of the mean.

On the other hand, neural tracking of several other characteristics was not modulated by attention. Neural tracking of musical note pitch value, for example, was similar when the music was attended (mean = 0.0087, S.D. = 0.0098) or ignored (mean = 0.0090, S.D. = 0.0078; p > 0.05). Pitch value tracking was greater in participants with more musical training (F = 5.05, p = 0.03). Figure 4 displays neural tracking of musical note pitch value across conditions. Neural tracking of melodic surprisal was also similar in the attended (mean = 0.0062, S.D. = 0.0086) and ignored (mean = 0.0060, S.D. = 0.0069; p > 0.05) conditions. There were no significant predictors of neural tracking of melodic surprisal, based on the model incorporating both long-term and short-term information. Figure 5 displays neural tracking of melodic surprisal across conditions. To follow up on these null findings, we used Bayesian paired samples t-tests in JASP software (35) to examine the extent to which the data provided evidence in support of the null hypothesis of equivalent tracking across attention conditions, using Bayes Factor interpretations proposed by Jeffreys (36). For melodic surprisal the BF_01_ was 6.27, indicating substantial support for the null hypothesis. For note value the BF_01_ was 6.23, indicating substantial support for the null hypothesis.

**Figure 4.**
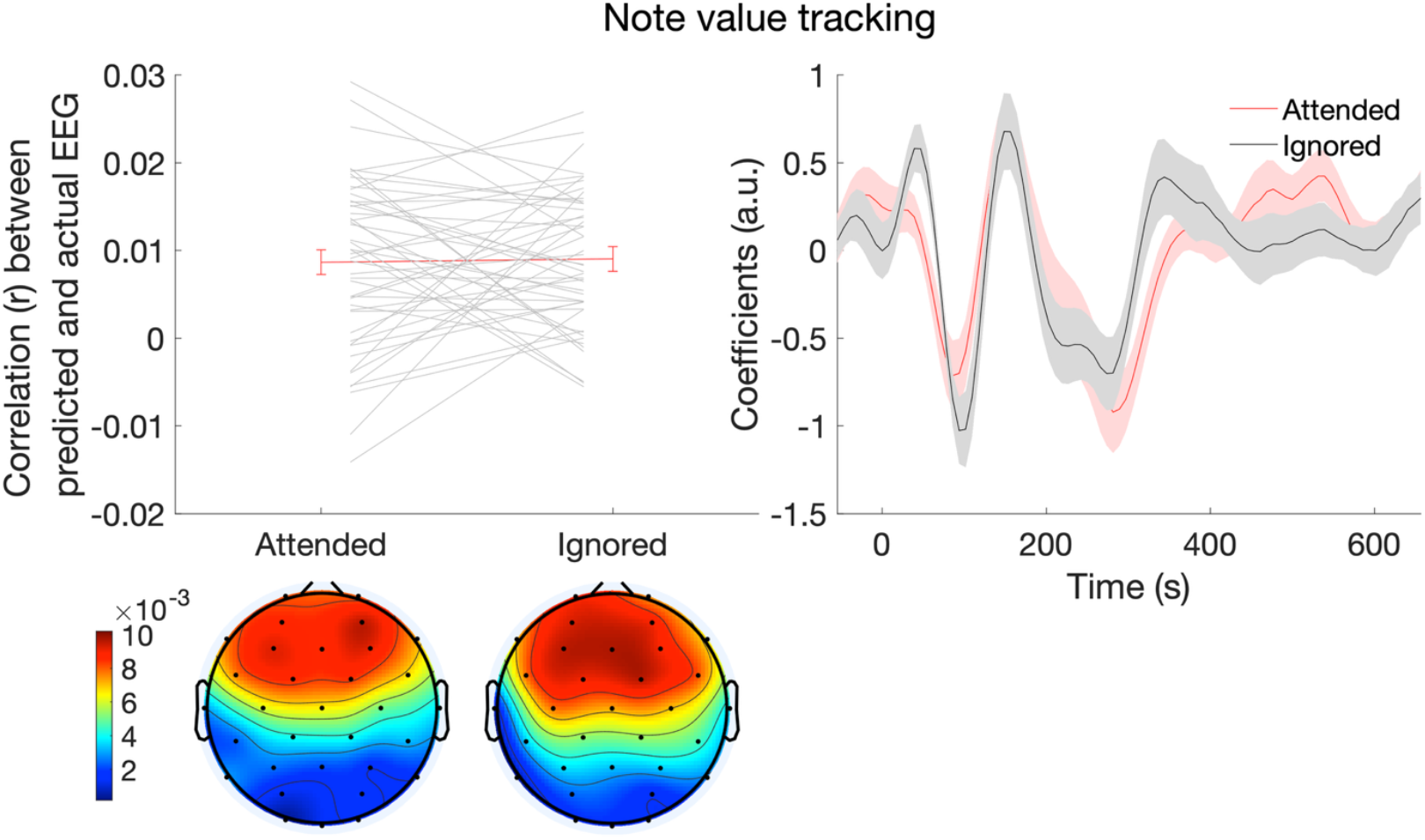
Effect of selective attention on musical note pitch value tracking. The outcome measure plotted on the left is the extent to which changes in the EEG can be predicted by a model making use of note value as a predictor. The right-hand side of the figure plots model coefficients across participants. Error bars display one standard error of the mean.

**Figure 5.**
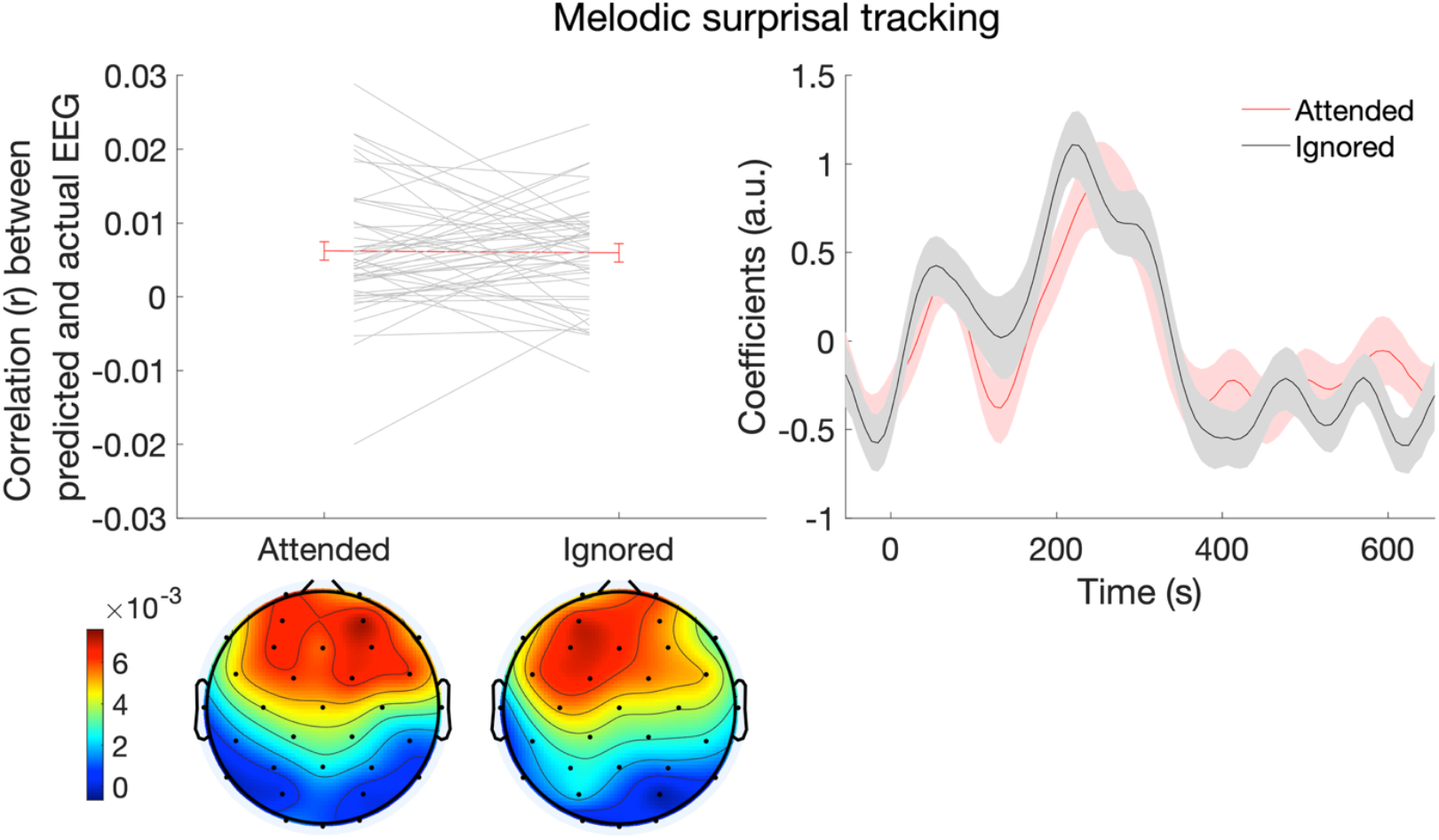
Effect of selective attention on melodic surprisal tracking. The outcome measure plotted on the left is the extent to which changes in the EEG can be predicted by a model making use of melodic surprisal as a predictor. The right-hand side of the figure plots model coefficients across participants. Error bars display one standard error of the mean.

One possible explanation for why melodic surprisal is not modulated by attention is that participants are able to make use of previously-learned information about the statistical characteristics of music to make predictions without drawing upon cognitive resources (22-23). To test this possibility, we investigated the effects of attention on melodic surprisal, calculated separately based on two IDyOM models: one incorporating long-term information only (trained on a large corpus of Western music but not tracking information about the current musical piece) and one incorporating short-term information only (trained exclusively and incrementally on the statistical characteristics of the current piece). Neural tracking of melodic surprisal based on the long-term model was unaffected by attention (p > 0.05): neural tracking was similar in the attended (mean = 0.0058, S.D. = 0.0081) versus ignored (mean = 0.0055, S.D. = 0.0082) conditions. A Bayesian paired samples t-test revealed a BF_01_ of 6.30, indicating substantial support for the null hypothesis. However, neural tracking of melodic surprisal based on the short-term model was greater in the attended condition (mean = 0.0058, S.D. = 0.0085) than in the ignored conditions (mean = 0.0022, S.D. = 0.0066; F = 4.98, p = 0.031). Moreover, although neural tracking was significantly greater than zero in the attended condition according to a Wilcoxon signed rank test (z = 4.03, p < 0.001), neural tracking was not significantly greater than zero in the ignored condition (z = 1.91, p > 0.05). Figures 6 and 7 display neural tracking of melodic surprisal based on the long-term and short-term models, respectively, across the attended and ignored conditions.

**Figure 6.**
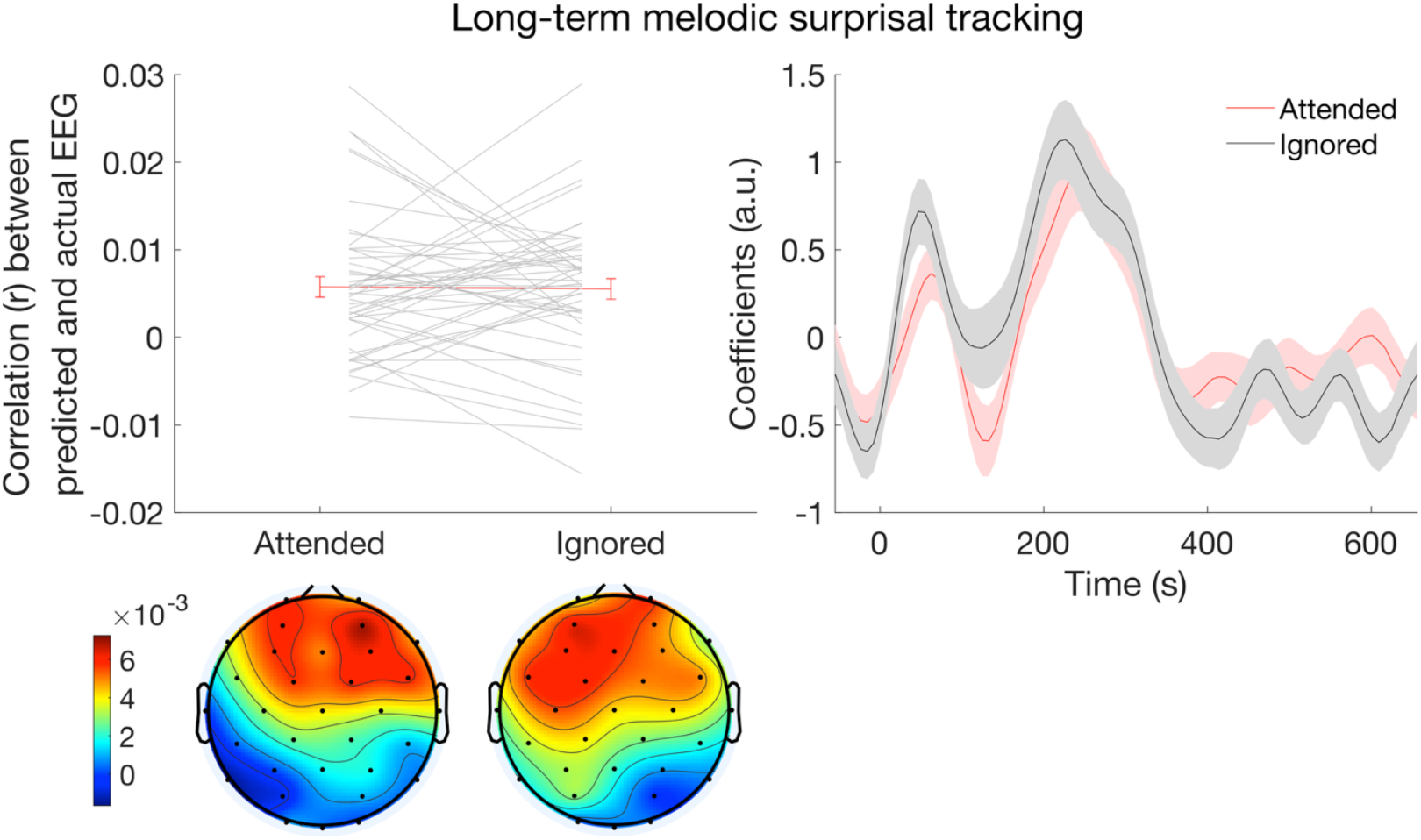
Effect of selective attention on melodic surprisal tracking based on a long-term model of musical prediction. The outcome measure plotted on the left is the extent to which changes in the EEG can be predicted by a model making use of melodic surprisal as a predictor. The right-hand side of the figure plots model coefficients across participants. Error bars display one standard error of the mean.

**Figure 7.**
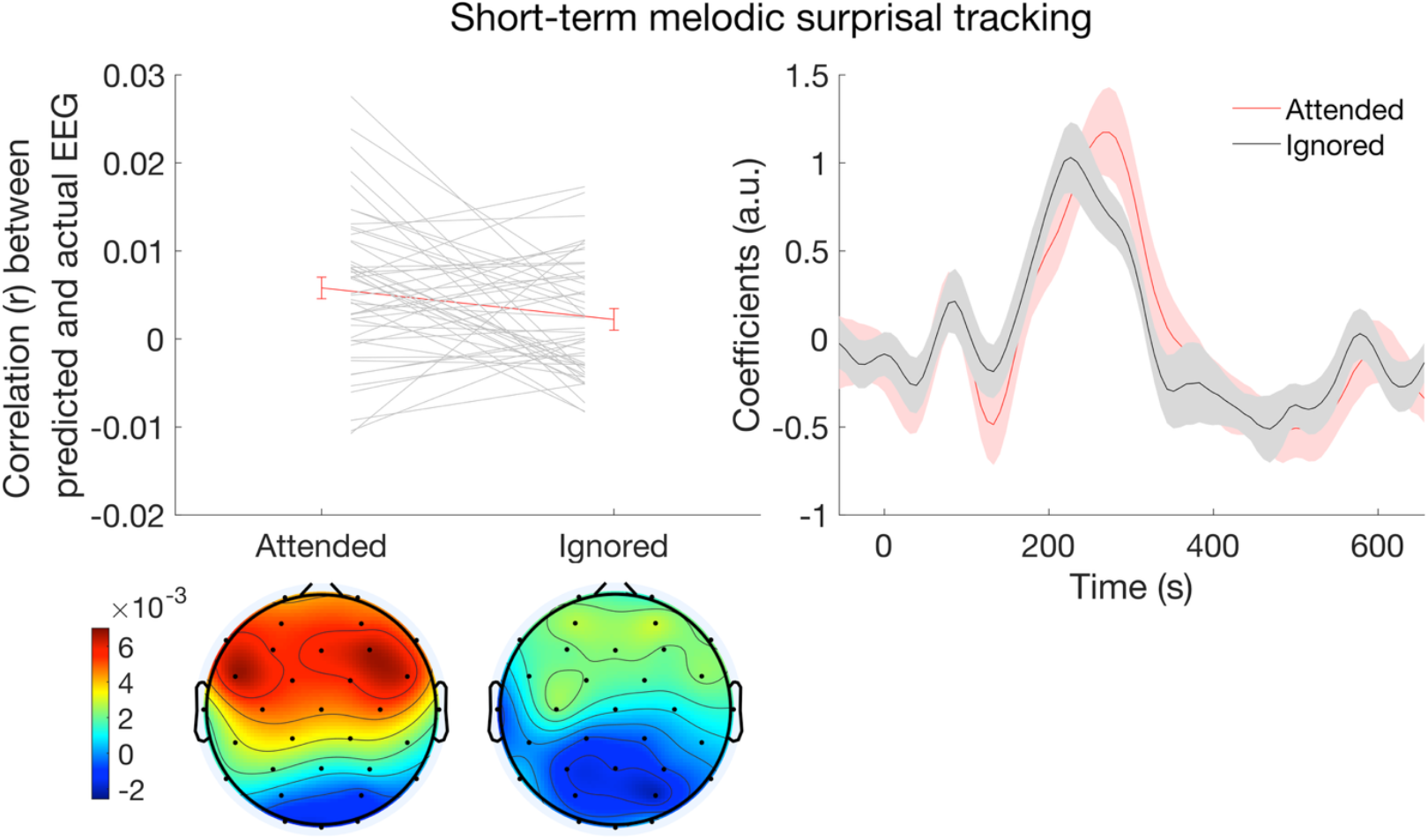
Effect of selective attention on melodic surprisal tracking based on a short-term model of musical prediction. The outcome measure plotted on the left is the extent to which changes in the EEG can be predicted by a model making use of melodic surprisal as a predictor. The right-hand side of the figure plots model coefficients across participants. Error bars display one standard error of the mean.

Next, we used frequency tagging analyses to investigate whether attention modulated neural tracking of the musical beat. First, we assessed normalized EEG magnitude at the frequency of the most common stimulus presentation rate. Stimulus rate tracking was greater in the attended condition (mean = 0.103, S.D. = 0.060) than in the ignore condition (mean = 0.083, std = 0.049; F = 10.2, p = 0.003), confirming that this analysis is sufficiently sensitive to detect effects of attention. We also found that stimulus rate tracking was greater in older participants (F = 6.0, p = 0.018) and in participants with more musical training (F = 11.1, p = 0.002), mirroring the findings for neural tracking of musical note onset. However, we found that neural tracking of the musical beat was no greater in the attention condition (mean = 0.011, S.D. = 0.028) than in the ignored condition (mean = 0.016, S.D. = 0.029; p > 0.05). We next used Bayesian paired sample t-tests to follow up on this null finding. The BF_01_ was 4.36, providing substantial support for the null hypothesis of no difference between attention conditions. There were no significant predictors of musical beat tracking. Figure 8 displays neural entrainment to the stimulus rate (left-hand side) and beat rate (right-hand side) across attention conditions.

**Figure 8.**
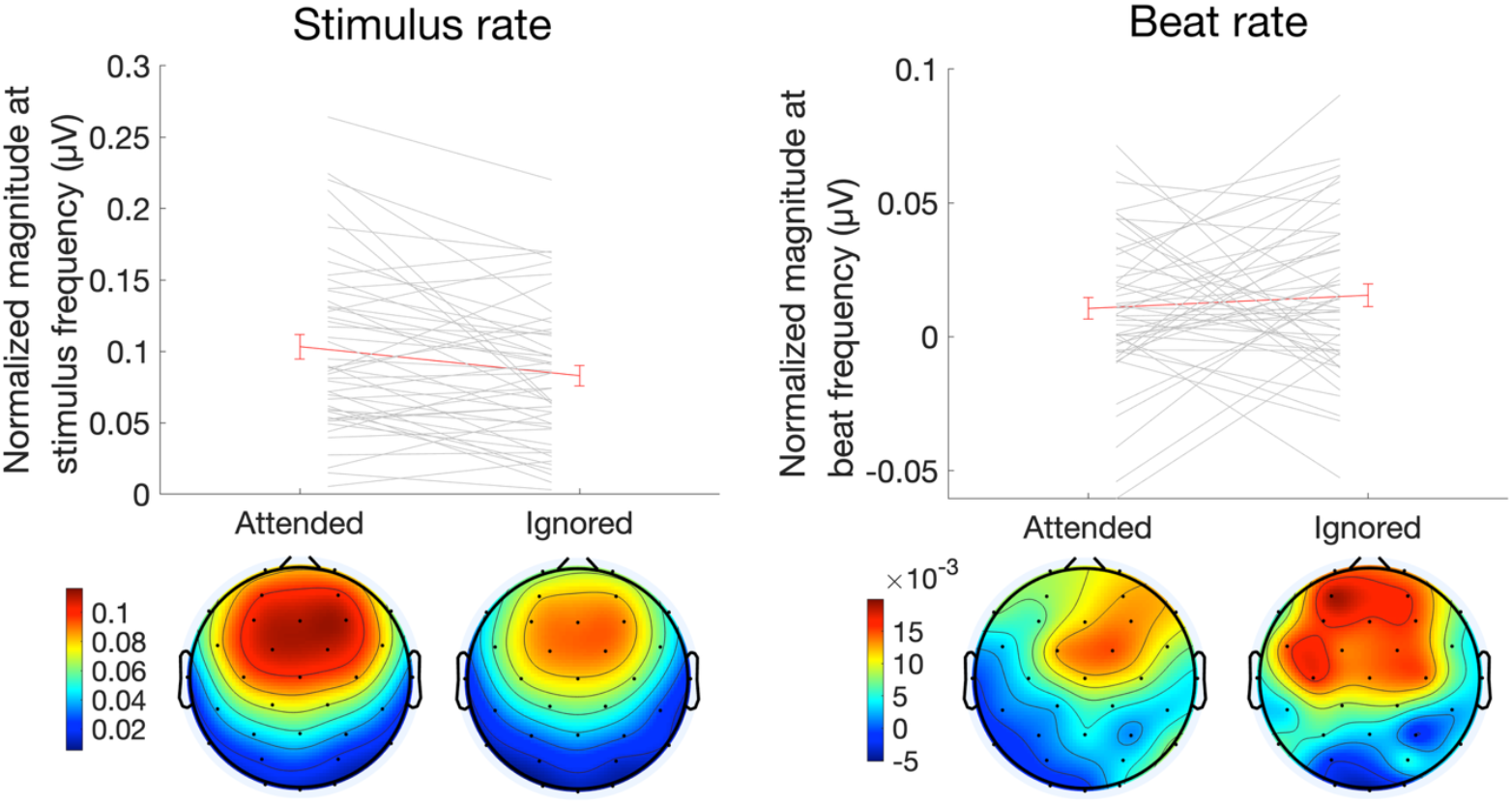
Effect of selective attention on stimulus rate (left) and musical beat (right) tracking. The outcome measure in both cases is magnitude at the frequency tagged to either the stimulus rate or musical beat rate, normalized by subtracting the surrounding 10 frequency values on either side of the critical rate (with 0.01 Hz resolution). Error bars display one standard error of the mean.

Finding no significant difference between attention conditions could theoretically be caused by a floor effect: if a feature of music is not tracked by the EEG, then there can be no effect of attention. To ensure that null effects of attention on tracking of particular features were not driven by a lack of neural entrainment to these features, we conducted follow-up 32-channel analyses for musical note pitch value, melodic surprisal, and musical beat. Using a Wilcoxon signed rank test, we found that r-values averaged across all channels and across both attention conditions were significantly different from zero for both pitch value (median = 0.0044, iqr = 0.0075, z = 5.3, p < 0.001) and melodic surprisal (median = 0.0036, iqr = 0.0071, z = 4.1, p < 0.001). Similarly, normalized EEG magnitude at the beat rate, averaged across channels and conditions, was significantly different from zero (median = 0.012, iqr = 0.025, z = 3.7, p < 0.001). To further confirm the success of neural tracking, we compared the shape of the TRF function across the two attention conditions, averaged across all participants and all 32 channels. The TRF shape was significantly correlated between attention conditions for both pitch value (r = 0.49, p < 0.001) and melodic surprisal (r = 0.90, p < 0.001).

Relationships between demographic predictors and neural tracking of stimulus features, collapsed across conditions, are illustrated in the following two figures. Figure 9 displays the relationship between age in years and neural tracking of speech amplitude and music note onset. Figure 10 displays the relationship between years of musical training and neural tracking of musical note onset and note pitch value.

**Figure 9.**
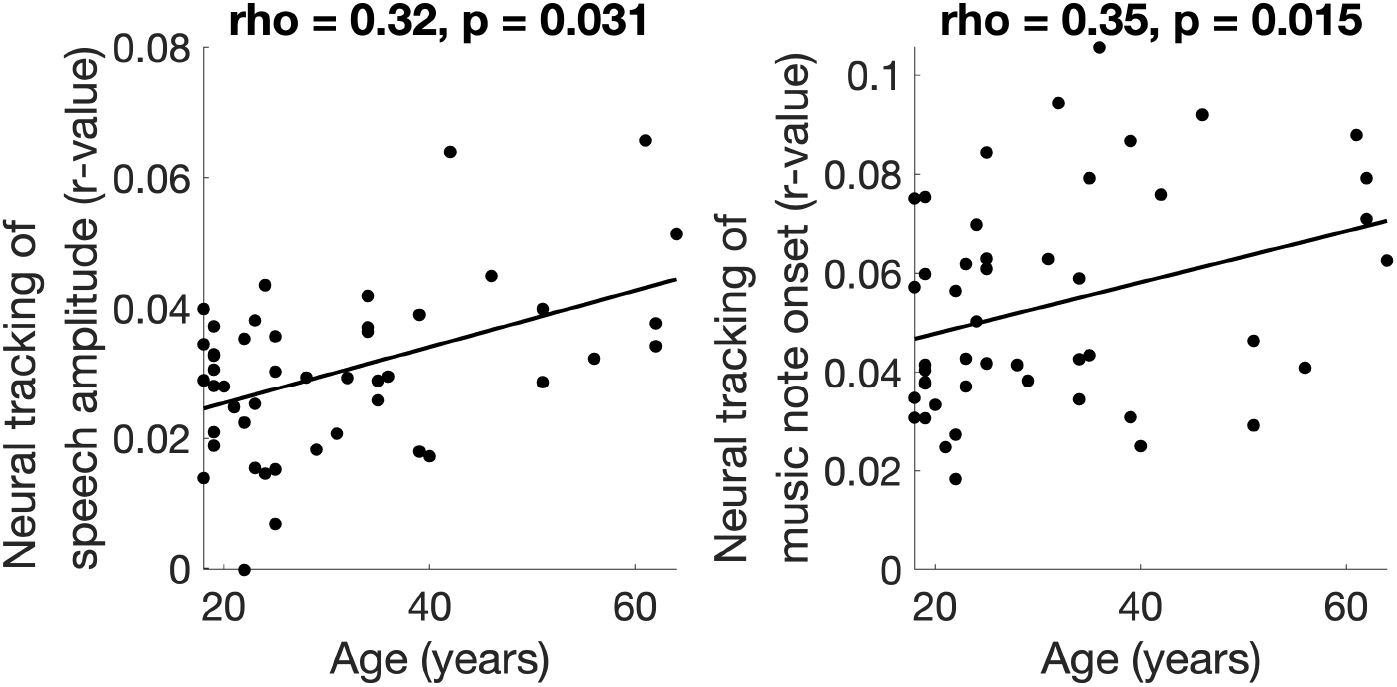
Scatterplots displaying the relationship between age in years and neural tracking of speech amplitude (left) and musical note onset (right).

**Figure 10.**
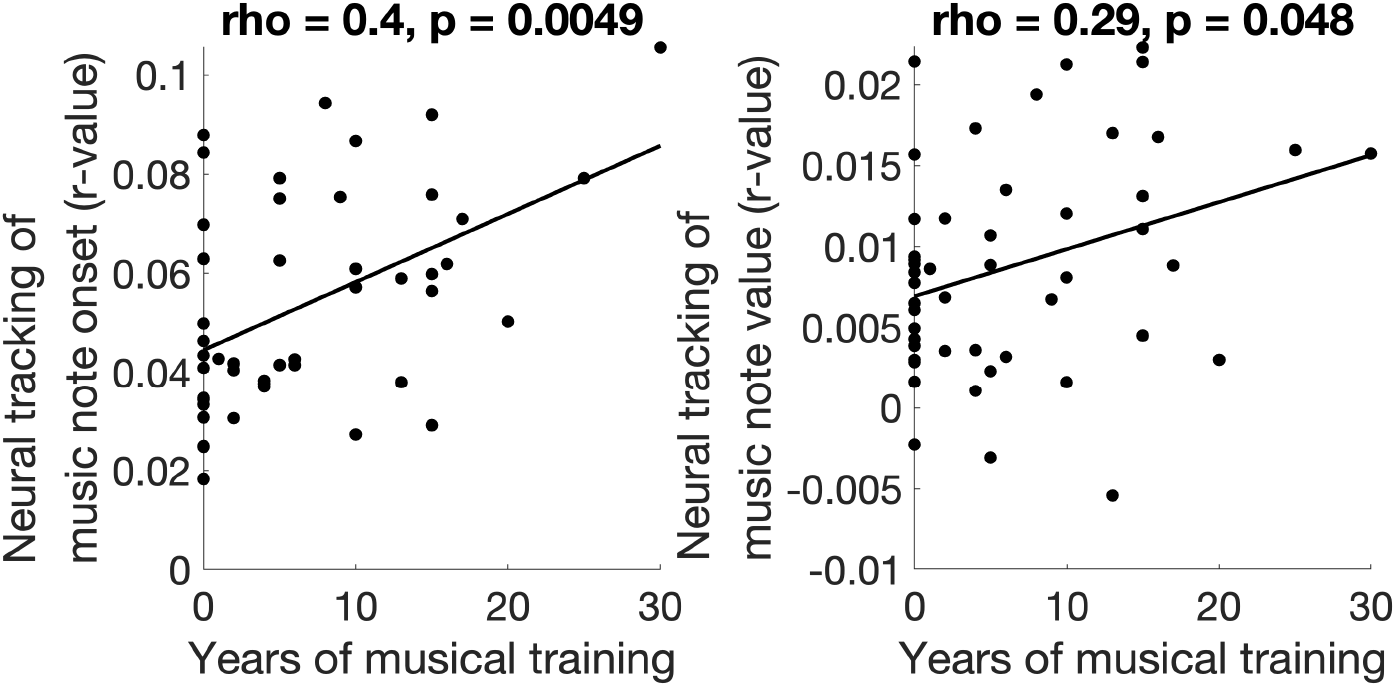
Scatterplots displaying the relationship between years of musical training and neural tracking of musical note onset (left) and music note value (right).

## Discussion

We asked how attention modulates neural encoding of musical features by simultaneously presenting participants with monophonic pieces of music by J. S. Bach in one ear and a single talker reading a Hemingway story in the other, asking them to attend to one or the other so as to perform a subsequent recognition memory test. We found that attention boosted encoding of acoustic features of music. Neural encoding of note onset, for example, was more robust when the music was attended. This finding replicates the results of prior research on selective attention to musical passages, which have shown that amplitude envelope encoding is enhanced by attention (37-41). We also found that attention enhanced encoding of the amplitude envelope of speech, which has been extensively demonstrated in prior research (42-47). Our results suggest that encoding of the amplitude envelope is the most important feature for reconstructing the direction of attention to auditory streams, even when these streams are drawn from different domains (48).

Effects of attention on music processing were not limited to encoding of the amplitude envelope, however: encoding of pitch interval size was more robust when stimuli were attended. Given that small intervals tend to predominate in music (11), encoding of pitch interval size is likely driven by responses to occasional pitches which suddenly diverge from the pitch range of the previous notes. This interpretation is supported by the shape of the TRF for encoding of pitch interval size, in which the largest component was similar in shape and timing to a P300, an ERP which is evoked by violations of short-term expectations (49). Our finding that pitch interval size encoding is enhanced by attention aligns, therefore, with prior findings that responses to violations of simple local expectations can be enhanced by attention. For example, when participants are presented with sets of identical tones, with changing frequencies between sets, the mis-match negativity response to the first tone in each set is enhanced when the stream is attended (17). The attentional enhancement of pitch interval size encoding fits the role of attention in predictive coding theory, in which attention up-weights the encoding of prediction errors, boosting learning (16).

In contrast to our finding that attention enhances encoding of acoustic features and low-level prediction in music, we found that encoding of high-level melodic prediction was unaffected by attention, as it was robust in both the attended and ignored conditions. That attention was limited to low-level processing is in stark contrast to prior research on selective attention to speech, which has found that attention effects are weaker for neural encoding of acoustic compared to linguistic features such as phonemic and semantic encoding (50-51). In fact, attention may be a requirement for processing certain features of speech: when two speech streams are presented to a participant simultaneously, lexical neural processing takes place only for the attended stream (52-53).

However, the results show that the null effect of attention holds only for effects of a long-term IDyOM model making static predictions based on training from a large corpus of melodies. This model simulates schematic effects on expectation based on an internal model of melodic structure derived through statistical learning during long-term exposure to music in a given style (Western tonal music in the present case both for participants and model). The lack of attentional effects on neural tracking of long-term prediction is consistent with evidence that early neural responses to stylistically unexpected chords (the Early Right Anterior Negativity or ERAN) are unaffected by attention (54). It has been proposed that schematic expectations are generated in an implicit and obligatory manner to account for why listeners continue to experience surprisal-induced pleasure even for music that is highly familiar (22-23). Hartung et al., (21) found that greater enjoyment was associated with better neural tracking of IDyOM’s melodic and rhythmic predictions while in behavioural research intermediate levels of predictability have been associated with greatest levels of pleasure (55-57).

By contrast, attentional affects were observed for neural tracking of predictions generated by the short-term IDyOM model, trained incrementally on each stimulus, simulating a psychological process of short-term learning for repeated patterns within a piece of music. Neural tracking of short-term prediction was greater in the attended than in the ignored condition, and in fact neural tracking of prediction in the ignored condition was not significantly different from zero. This suggests that attention is required for local melodic learning: extracting patterns from the current listening episode and using them to guide expectations. Evidence that pattern learning for immediate repetitions of patterns in rapid tone sequences over a span of a few seconds can operate without direction of attention (58; 59) suggests that the attentional effects in the present experiment may be due to the length of time over which patterns reoccur within the stimulus or the complexity of the patterns and approximate nature of the repetition. Since learning and prediction both occur within the experimental session, it is not possible to distinguish whether attention is required for learning, prediction or both.

We also found that neural tracking of the musical beat was unaffected by attention. Given that musical beats can be seen as a generative model used to predict note timing (25-26), this again suggests that high-level musical prediction does not require attention but can take place even when complex music is ignored. Prior research on the effects of attention on musical beat tracking is somewhat mixed. Research suggests that musical beats can speed up performance on unrelated tasks such as visual target detection and visual word recognition (60-62). Moreover, pupil dilation responses and ERPs to deviant sound events and omissions are modulated by musical beat structure even when attention is directed to an alternate task (63-64). However, directing attention away from complex auditory rhythms and towards visual stimuli was found to change the pattern of fMRI activation, including decreased activity in the basal ganglia, supplementary motor area, and pre-motor cortex (65). Neural encoding of the beat of auditory rhythms has also been shown to be enhanced by attention to the rhythms, relative to a visual task (29, 66), although another study found that cross-domain selective attention had no effect on musical beat encoding (28). One possible explanation for our finding that musical beat encoding was robust even when the music was ignored is that our passages were relatively long, rhythmically simple and consistent, possibly enabling participants to quickly figure out the beat timing and then implicitly maintain it while directing attention elsewhere.

We found that the effects of attention on neural encoding of musical features were not modulated by musical expertise. However, we found that neural encoding of note onset and note pitch value was enhanced in participants with more musical training across both the attended and ignored conditions. This suggests that musical training does not enhance the ability to attend to music, and that the effects of musical training on processing of musical features take effect at a pre-attentive level. This finding conflicts somewhat with a previous report that musicians are better able to perceive music embedded in informational maskers consisting of multiple streams of music (31). It is possible that musicians are not better at directing attention to music, but instead at performing stream segregation to separate out musical streams, prior to stream selection. Our finding of enhanced encoding of note onset aligns with previous reports of greater cortical tracking of the acoustic envelope of music in musicians compared to non-musicians (32). However, here, by simultaneously recording neural responses to speech, we were able to show that the effects of musical training are domain-specific, boosting neural encoding of the amplitude envelope of music but not speech. This finding is consistent with a recent report that musicians do not show enhanced neural encoding of speech in noise (67). Our finding that encoding of melodic surprise is not enhanced in participants with more musical training is in line with previous reports of equivalent encoding of melodic prediction error in ecologically valid music between musicians and non-musicians (34). However, it conflicts somewhat with the report that mis-match responses to deviant notes in five-note melodies are greater in musicians compared to non-musicians (33). One possibility is that musicians may be better able to make melodic predictions when given very little context, as in isolated five-note melodies, while the increased context and repetition present in streams of ecologically valid music may boost prediction in non-musicians sufficiently that they achieve the same level of prediction as musicians.

We found that aging was linked to enhanced encoding of both speech amplitude and note onset. This aligns with prior research showing greater neural entrainment to speech amplitude envelope between younger/middle and older adulthood (68-69) and extends this finding to music. The mechanism underlying this effect, therefore, must be domain-general. One possibility is that cortical auditory responses are boosted to compensate for less reliable subcortical sound representation. Another possibility is that this increased encoding reflects listening effort, as enhanced encoding of speech amplitude has also been found when individuals listen to speech in a second language relative to a first language (70). Aging, however, was not linked to changes in the modulation of neural tracking of music or speech by attention. The ability to selectively attend to music, therefore, does not decline with age within the age range tested here (18 to 64 years).

One limitation of this study is that we used monophonic musical stimuli (music in which there is only a single note playing at a time). This choice simplified the calculation of melodic surprisal, but it prevented us from investigating neural encoding of certain musical features, such as harmony. EEG responses to surprising chords, based on the preceding harmonic context, occur as early as 200 milliseconds (71) and as late as 600 milliseconds (72) after chord onset. It remains unclear, however, whether a musical sequence must be attended for harmonic prediction to occur. Prior studies of the effects of selective attention on EEG responses to unexpected chords have found conflicting results, with one study reporting equal responses to harmonic surprise regardless of attention (73), one study reporting an effect of attention on early responses (74), and a third study reporting an effect of attention only on late responses (75). Moreover, these studies used short, simple chord sequences, so the effects of attention on harmonic prediction in ecologically valid music listening remain unknown. Based on the results of the current paper, we predict that harmonic surprisal encoding in ecologically valid music will be robust even when attention is directed away from the music and towards a concurrent speech stimulus. However, we predict that predictions based on a short-term model of musical expectation will be better tracked in the attended condition.

In conclusion, we find that selective attention boosts neural encoding of low-level acoustic features of music, but that high-level musical prediction is robust even when music is ignored. Thus, surprisingly sophisticated music listening can be carried out even when music fades into the background. This automaticity of music listening, possibly due to the repetitive nature of music compared to other art forms, may be what enables it to play such an effective supporting role in multimedia art.

## Methods

### Participants

50 participants were recruited from the greater London area. Data from 2 participants were not analyzed because they performed below chance on the attention task (see below for more details). Data from 48 participants (31 female) were analyzed. Participants were all native speakers of English. They reported a mean age of 31.3 years (standard deviation 13.6, range 18 to 64) and a mean of 6.6 years of musical training (S.D. = 7.5, range 0 to 30). Informed consent was obtained from all participants. To reimburse participants for their time, they were given either course credit or £10 per hour. Experiment procedures were approved by the ethics committee of the School of Psychological Sciences at Birkbeck, University of London.

### Stimuli

20 total audio tracks were created. Each of these tracks featured speech in one ear and music in the other. The music stimuli were a series of short monophonic Bach pieces, extracts of approximately 150 seconds from works for solo violin and flute, encoded into MIDI format and synthesized using a piano timbre. (These pieces were retrieved in .midi form from jsbach.net.) A few of the music pieces were slightly edited to be monophonic (in a few instances where multiple notes appeared at once, all but one of the notes was removed). The speech stimuli consisted of passages from the Hemingway story The Old Man and the Sea. This recording has been used previously to investigate neural tracking of the amplitude envelope and pitch contour of speech (76). The speech was divided into blocks a few minutes long, matching the duration of the music pieces. The root mean square amplitude of the speech and music stimuli were equated. Stimuli were presented at max 80 dB SPL at a sampling rate of 44,100 Hz using MATLAB (77) via 3-M E-A-RTONE 3-A insert earphones.

### Procedure

For half of the stimuli, participants were asked to attend to the music. For the rest of the stimuli, participants were asked to attend to the speech. The attention condition was blocked; in other words, all the attend speech trials were presented together, as were all the attend music trials. At the beginning of each block, participants heard recorded instructions telling them to listen to either the speech or the music and to ignore the competing stimulus. Two seconds after the end of each trial, participants were presented with a 5-second clip of either music (in the attention to music condition) or speech (in the attention to speech condition). For half of the trials, this clip was an excerpt of the stimulus the participant had just heard; in the other half, it was taken from one of the other stimuli. Assignment of stimuli to these two conditions was randomized. The task was to indicate whether the clip was part of the recording presented in the just completed trial. Participants responded by typing “yes” or “no” on a keyboard.

Condition order (attend speech versus attend music) was counter-balanced across participants. Assignment of the stimuli to the two attention conditions was randomized for each participant. For half of the stimuli in each attention condition, speech was in the left ear while music was in the right ear, while for the remaining half, music was in the left ear while speech was in the right; assignment of stimuli to these two laterality conditions was randomized. Stimulus order was also randomized separately for each participant.

### EEG data collection

EEG data was collected via a BioSemi ActiveTwo system with a sample rate of 16,384 Hz and digitized with 24-bit resolution. 32 electrodes embedded in a cloth cap were arranged according to the 10-20 system, with reference electrodes attached at the earlobes and CMR/DRL electrodes functioning as ground. Electrode impedance was maintained below 20 kΩ throughout the session. An RTBox (78) sent triggers to the EEG data collection computer and detected the onset of the auditory stimulus on each trial so that EEG data could be precisely aligned with stimulus timing.

### EEG data processing

All EEG data processing and analysis were carried out in MATLAB (77) using the FieldTrip M/EEG analysis toolbox (79) in combination with in-house scripts. EEG data were downsampled to 128 Hz then re-referenced offline to the average of the two earlobe electrodes. Data were low-pass filtered below 15 Hz using ft_processing.m with default settings (a sixth-order Butterworth filter), then high-pass filtered above 1 Hz with a fourth-order filter. ICA was then used (ft_componentanalysis, “runica” method) to identify components whose topography and time course suggested the influence of eye blinks and eye movements; these components were removed using ft_rejectcomponent. Finally, EEG data were separated into 20 epochs aligned with the onset and offset of each auditory stimulus.

### Stimulus feature extraction

The amplitude envelope of the speech was extracted using the Hilbert transform. Characteristics of the musical stimuli were extracted by consulting the .midi data and then aligning the resulting values with the sound file, downsampled to 128 Hz. First, note onsets were marked at the beginning of each note. Additional characteristics were then aligned with the note onset vector, consisting of non-zero values at note onset and zeros elsewhere. These characteristics included note pitch value (in semitones from 440 Hz) and absolute pitch interval size (the absolute difference between the pitch of each note and the previous note in semitones). Next, melodic prediction was calculated using the IDyOM (Information Dynamics of Music; 7) model. This model predicts the next note of a melody based on pitch and timing information. The model incorporates information based both on patterns across a large corpus of music (long-term) and patterns within a single piece of music (short-term). Melodic surprisal was calculated as the information content of each note, as provided by IDyOM. This is the extent to which a note is surprising, given the preceding melodic context. Melodic surprisal was calculated based on long-term and short-term models, as well as an overall model incorporating both long-term and short-term information. Note pitch value, absolute pitch interval size, and melodic surprisal were z-scored prior to analysis.

### Multivariate temporal response function analysis

Neural tracking of speech and music was assessed using the multivariate temporal response function (mTRF, 80) implemented in Matlab. This technique uses regularized regression to build a model relating characteristics of continuous naturalistic stimuli to changes in the EEG signal at a range of time lags. We used a doubly-nested cross-validation procedure to identify the regularization parameter (lambda) from a set of potential values (10^-6^, 10^-4^, 10^-2^, 1, 10^2^, and 10^4^) and then test the model. First, we divided the ten trials in each condition into nine training trials and one test trial. For each potential lambda value, the model was trained on eight of the nine training trials and tested on the remaining trial by correlating the predicted with the actual EEG, and this process was repeated so that each trial was selected as the testing trial once. The lambda was then chosen which maximized the EEG prediction across trials and channels. Finally, this lambda was used to train the model across all nine training trials and this model was tested on the remaining trial. This procedure was then repeated ten times, with each trial serving as the test data once. The prediction accuracy (r value) and model coefficients were averaged across all four models.

The time lags chosen to train the model varied across stimulus features. For acoustic features of the speech and music, including amplitude envelope and note onset, we trained the model on lags ranging from -50 to 350 ms. This time region was chosen to match previous research on neural tracking of acoustic features of speech (76, 81) and music (24). For note pitch value, pitch interval size, and melodic surprisal, the analysis window ranged from -50 to 650 ms, matching previous research on neural tracking of melodic features of music (Kern et al. 2022).

When analyzing neural encoding of note pitch value, pitch interval size, and melodic surprisal, we first controlled for neural encoding of note onset. This ensured that we specifically measured neural tracking of these musical features, without any confounding influence of note onset. To do this we first trained a model to predict the EEG from note onset timing. We then subtracted the predicted EEG from the actual to extract model residuals. These residuals were then entered into the next round of mTRF analysis, to be predicted by the musical features.

### Frequency tagging musical beat analysis

Neural tracking of the beat of music was assessed using frequency tagging. This approach was made possible by our use of isochronous midi-synthesized music. For each of the 20 musical pieces, the stimulation rate and beat rate were assessed. The stimulation rate was set to the rate corresponding to the most common inter-stimulus interval throughout the piece. The beat rate was calculated manually by the authors by listening to each piece and determining the rate at which they most naturally synchronized when tapping to the beat. (The beat rate could not be assessed for one of the pieces, due to the tempo being much slower than that of the other pieces; this piece was removed from the analysis.)

For each epoch of the EEG, corresponding to the entire period during which one of the pieces was presented, a fast Fourier transform was used to extract the spectrum, with a resolution of 0.01 Hz. The normalized magnitude at the target frequencies was then calculated by assessing the magnitude at the frequencies equivalent to the stimulation and beat rates and then subtracting from these values the average magnitude of the surrounding 20 frequencies (10 on each side, corresponding to 0.1 Hz below to 0.1 Hz above the target frequency). This process was carried out for each channel. We then calculated the nine channels with the highest degree of frequency tagging, averaged across both target frequencies (stimulation rate and beat rate), both attention conditions, and all stimuli. Finally, stimulation rate and beat rate tracking were averaged across these channels and across stimuli separately for the attend music and attend speech conditions.

### Statistical analysis

For the mTRF analyses, overall tracking of each characteristic was compared between the attend music and attend speech conditions. This was done by converting the correlation between predicted and actual EEG to z-scores using Fischer’s r to z conversion, then collapsing across the 9 channels with the highest tracking across both attention conditions. A mixed model (using lmer in R) was then used to compare neural tracking between the attended and unattended conditions, with years of musical training and age as additional between-subjects factors. The regression equation was: tracking ∼ attention * musical_training * age + (1 | ID). This analysis was carried out for speech amplitude, musical note onset, musical note pitch value, pitch interval size, and melodic surprisal. For the beat tracking analyses, normalized EEG magnitude at the stimulation and beat rates was compared between the attended and unattended conditions. Similar to the mTRF analyses, this was done using mixed models in R, with the same regression equation (tracking ∼ attention * musical_training * age + (1 | ID)). Null effects of attention were followed up on by using Bayesian t-tests conducted in JASP software (35) to examine the strength of evidence for equivalence between the two attention conditions. For any feature whose tracking was not modulated by attention, we additionally ran follow-up analyses across all 32 channels to test whether there was significant neural tracking, to make sure that null effects did not simply reflect an overall lack of neural entrainment to these features.

The shape of the TRF models was compared across conditions. This was done by extracting the model weights for each time point, collapsed across channels, and then comparing between conditions using a paired t-test. The resulting p-values were corrected for multiple comparisons across time points using cluster analysis (83), with a threshold for cluster inclusion of p < 0.05, an alpha of 0.05, and 1000 iterations. However, no differences between conditions survived correction for multiple comparisons for any of the stimulus features, so these analyses are not reported on further.

## Data availability

Data, analysis scripts, and stimuli are available at https://osf.io/ed2rf/.

## Notes

https://osf.io/ed2rf/

### Competing Interest Statement

The authors have declared no competing interest.

